# Oxymatrine Induces Ferroptosis in MKN28 Gastric Cancer Cells

**DOI:** 10.1101/2025.11.22.689973

**Authors:** Liqiong Ma, Ruiqing Mian

## Abstract

This study aimed to investigate the effect and potential mechanism of oxymatrine (OMT) on MKN28 gastric cancercells. Through a series of experiments including CCK8 cell viability assay, lipid peroxidation level detection, and Fe^2+^ concentration measurement, it was found that OMT significantly inhibited the viability of MKN28 cells in a dose- and time-dependent manner. OMT could induce an increase in intracellular lipid peroxidation level and iron ion concentration, while activating ferroptosis-related signaling pathways, indicating that OMT can induce ferroptosis in MKN28 cells. This study provides a new theoretical basis and potential target for the application of OMT in gastric cancer treatment.

## 1. Introduction

Gastric cancer is one of the malignant tumors with high incidence and mortality worldwide ^[1]^. Despite the continuous development of current therapeutic approaches such as surgery, chemotherapy, and radiotherapy, the prognosis of patients remains unsatisfactory, especially for those with advanced gastric cancer. Thus, there is an urgent need to explore new effective therapeutic methods and drugs. Ferroptosis is a novel form of programmed cell death characterized by iron-dependent lipid peroxidation, which is significantly different from traditional cell death patterns such as apoptosis and necrosis ^[2]^. In recent years, a growing number of studies have demonstrated that inducing ferroptosis in tumor cells provides a new strategy and direction for cancer therapy ^[3]^. Oxymatrine (OMT) is an alkaloid extracted from the leguminous plants Sophora flavescens Ait. or Sophora alopecuroides L., which exhibits multiple biological activities including anti-inflammatory, antiviral, and antitumor effects. Previous studies have shown that OMT can inhibit proliferation and induce apoptosis in various tumor cells ^[4]^; however, whether OMT can induce ferroptosis in MKN28 cells has not been reported. Therefore, this study aims to investigate the effect of OMT on MKN28 cells, clarify whether it can induce ferroptosis in these cells, and provide a theoretical basis for the application of OMT in gastric cancer treatment.

## 2. Materials and Methods

### 2.1 Reagents

Oxymatrine (Shanghai Aladdin Biochemical Technology Co., Ltd.); RPMI 1640 cell culture medium (Gibco); penicillin-streptomycin double antibody (Gibco); CCK-8 kit (Shanghai Yeasen Biotechnology Co., Ltd.); iron ion detection kit (Nanjing Jiancheng Bioengineering Institute); protein extraction kit (Beyotime Biotechnology Co., Ltd.); GPX4, SLC7A11, and Nrf2 antibodies were purchased from Proteintech Group, Inc.

### 2.2 Cell Culture

Gastric cancer MKN28 cells were cultured in RPMI 1640 medium supplemented with 10% FBS and 1% penicillin-streptomycin double antibody in an incubator at 37°C with 5% CO_2_. Subsequent experiments were performed when the cells reached the logarithmic growth phase.

### 2.3 CCK-8 Assay for Cell Viability

MKN28 cells were seeded into 96-well plates at a density of 5×10^3^ cells per well. After 24 h of culture, different concentrations (0, 4, 6, 8 mg/mL) of OMT were added, and the cells were further cultured for 24 h or 48 h. Then, 10 μL of CCK-8 reagent was added to each well. After incubation for 2 h, the absorbance (OD value) was measured at 450 nm using a microplate reader, and cell viability was calculated. Cell viability (%) = [(OD value of experimental group - OD value of blank group) / (OD value of control group - OD value of blank group)] × 100%.

### 2.4 Determination of Lipid Peroxidation Level

MKN28 cells were seeded into 6-well plates at a density of 1×10^6^ cells per well. After 24 h of culture, OMT was added, and the cells were cultured for another 24 h. The intracellular malondialdehyde (MDA) content was determined according to the instructions of the lipid peroxidation detection kit to reflect the level of intracellular lipid peroxidation.

### 2.5 Detection of Fe^2+^ Concentration

Total protein was extracted with a commercial protein extraction kit, and protein concentration was quantified using the BCA assay. Equal amounts of protein samples were separated by SDS-PAGE and transferred onto PVDF membranes. Membranes were blocked with 5% non-fat milk for 2 h, then incubated with primary antibodies (1:1000 dilution) at 4°C overnight. After TBST washes, membranes were incubated with secondary antibodies (1:5000 dilution) at room temperature for 2 h. Following additional TBST washes, protein bands were visualized using an enhanced chemiluminescence (ECL) kit. Band gray values were analyzed with ImageJ software, and relative target protein expression was normalized to β-actin as the internal reference.

### 2.7 Statistical Analysis

All experiments were repeated three times, and the data were expressed as mean ± standard deviation 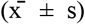. Statistical analysis was performed using GraphPad Prism 8.0 software. Comparisons between groups were conducted by one-way analysis of variance (One-way ANOVA), and a p-value < 0.05 was considered statistically significant.

## 3. Results

### 3.1 Effect of Oxymatrine on the Viability of MKN28 Cells

The results of the CCK-8 assay showed that compared with the control group, treatment of gastric cancer MKN28 cells with different concentrations (0, 4, 6, 8 mg/mL) of oxymatrine (OMT) for 24 h or 48 h significantly reduced cell viability in a dose- and time-dependent manner (*p* < 0.05). These findings indicate that OMT can effectively inhibit the proliferation of gastric cancer MKN28 cells.

### 3.2 Effect of Oxymatrine on the Lipid Peroxidation Level of MKN28 Cells

The results of lipid peroxidation detection indicated that compared with the control group, treatment of MKN28 cells with OMT for 24 h significantly increased intracellular malondialdehyde (MDA) content, while significantly decreasing GSH and SOD levels (*p*<0.05). These results suggest that OMT can induce lipid peroxidation in gastric cancer MKN28 cells (Figure 2).

**Figure 1.**
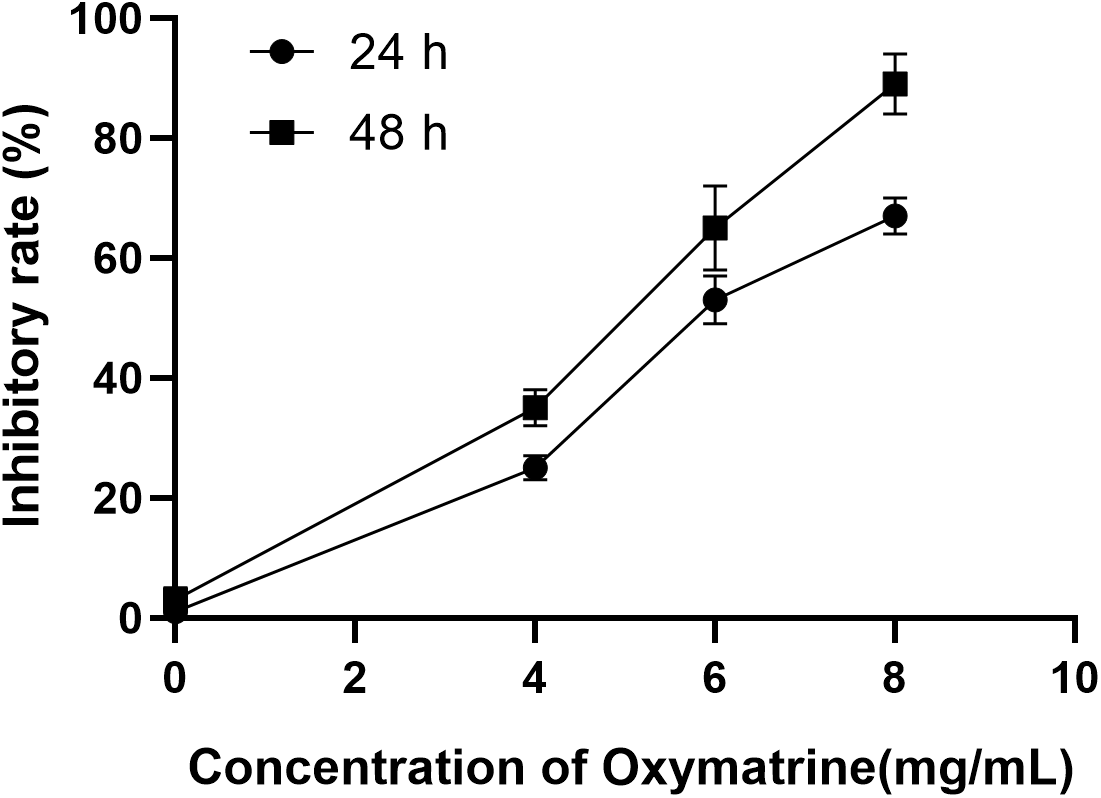
Effect of oxymatrine on the viability of MKN28 cells. The CCK-8 assay shows the changes in cell viability of each group after treatment with different concentrations (0, 4, 6, 8 mg/mL) of OMT for 24 h or 48 h.

**Figure 2.**
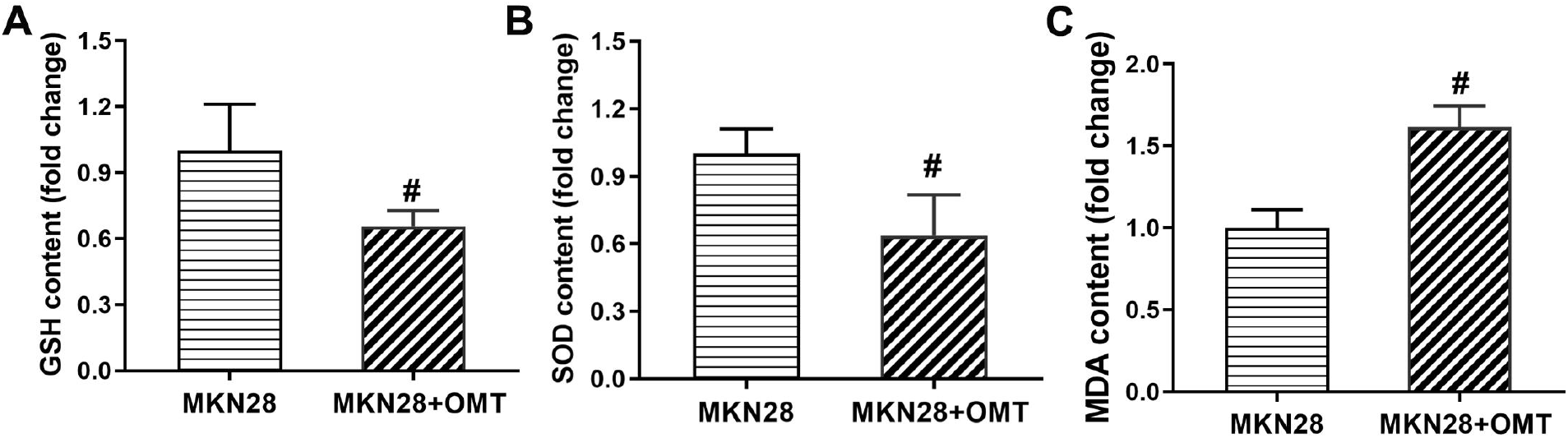
Effect of oxymatrine on the lipid peroxidation level of gastric cancer MKN28 cells. Changes in MDA, GSH, and SOD levels in each group. *p* < 0.05.

### 3.3 Effect of Oxymatrine on Fe^2+^ Concentration of MKN28 Cells

The results of Fe^2+^ detection showed that compared with the control group, treatment of gastric cancer MKN28 cells with OMT for 24 h significantly increased intracellular Fe^2+^ concentration (*P*< 0.05). These results indicate that OMT can promote Fe^2+^ accumulation in gastric cancer MKN28 cells (Figure 3).

**Figure 3.**
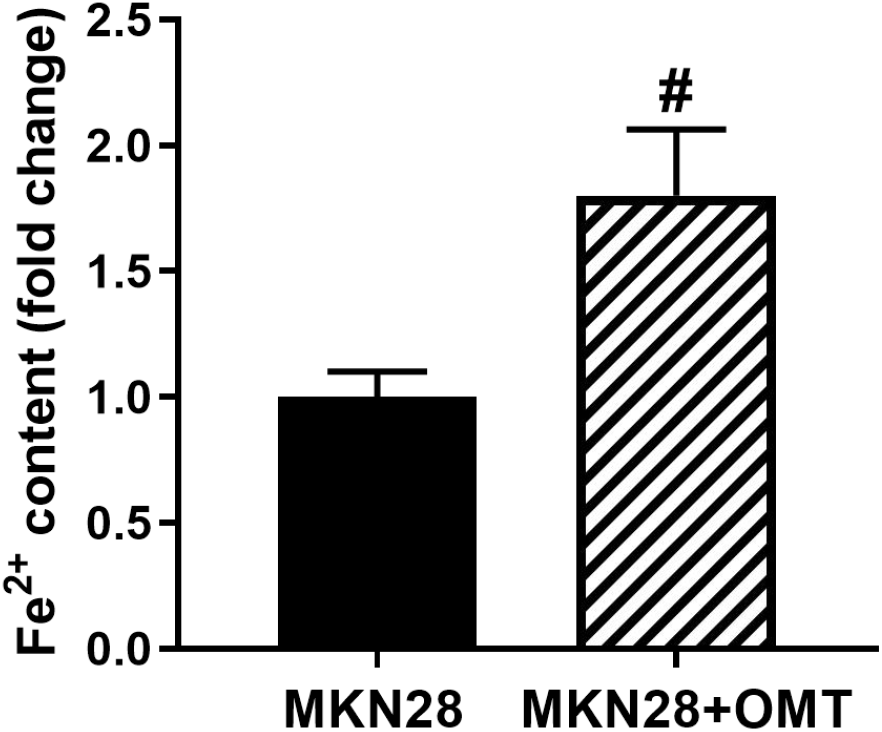
Effect of oxymatrine on the Fe^2+^ concentration of MKN28 cells. Changes in Fe^2+^ concentration in each group. *p* < 0.05.

### 3.4 Effect of Oxymatrine on the Expression of Ferroptosis-Related Proteins in MKN28 Cells

Western blot results showed that compared with the control group, treatment of gastric cancer MKN28 cells with OMT for 24 h significantly increased the expression of ACSL4 protein (*p*< 0.05) and significantly decreased the expressions of GPX4 and SLC7A11 proteins (*p*< 0.05). Consistent with the gene expression results, these findings further demonstrate that OMT can regulate the expression of ferroptosis-related proteins (Figure 4).

**Figure 4.**
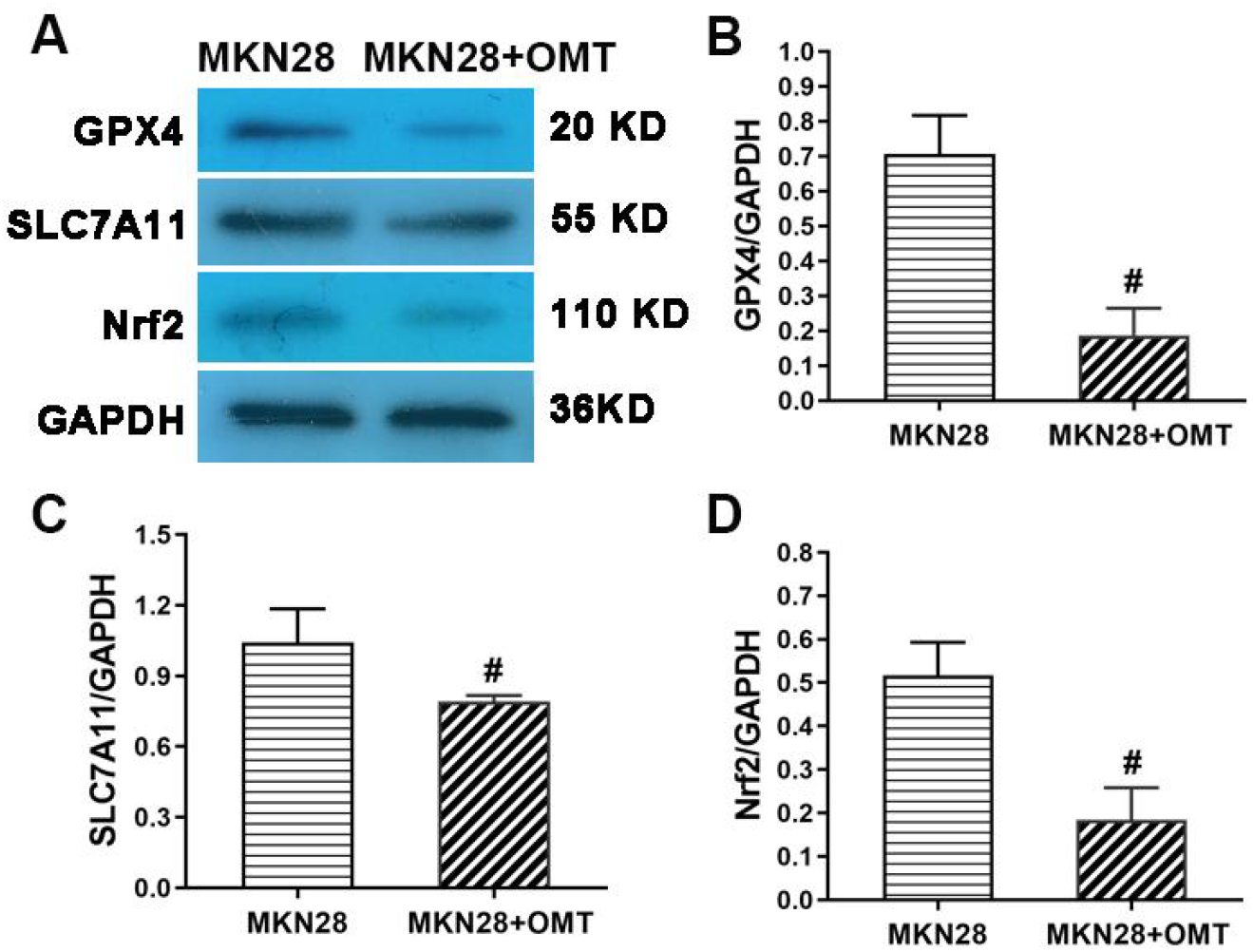
Effect of oxymatrine on the expression of ferroptosis-related proteins in MKN28 cells. Western blot analysis of ACSL4, GPX4, and SLC7A11 protein expressions in each group.

## 4 Discussion

This study is the first to demonstrate that oxymatrine (OMT) can induce ferroptosis in MKN28 cells. The results showed that OMT significantly inhibited the proliferation of MKN28 cells in a dose- and time-dependent manner, suggesting a potent cytotoxic effect of OMT on these cells. Lipid peroxidation is one of the key characteristics of ferroptosis. In the present study, OMT treatment significantly increased the intracellular malondialdehyde (MDA) content in gastric cancer MKN28 cells, indicating the occurrence of intracellular lipid peroxidation, which provides crucial evidence for OMT-induced ferroptosis. The initiation of ferroptosis relies on intracellular iron ion accumulation. Our findings revealed that OMT could enhance the intracellular iron ion concentration in MKN28 cells, further supporting the notion that OMT induces ferroptosis.

Lipid peroxidation stands as a hallmark biochemical feature that distinguishes ferroptosis from other forms of programmed cell death, such as apoptosis or necroptosis ^[5]^. In the present work, OMT treatment triggered a significant elevation in intracellular malondialdehyde (MDA) levels in MKN28 cells (Figure 2), a well-validated marker of lipid peroxidation that reflects the extent of oxidative damage to membrane polyunsaturated fatty acids (PUFAs). This observation is particularly noteworthy, as MDA accumulation directly correlates with the breakdown of cellular redox homeostasis—a key prerequisite for ferroptosis initiation. Concomitantly, OMT treatment was associated with a reduction in glutathione (GSH) levels and superoxide dismutase (SOD) activity (Figure 2), further supporting the notion that OMT disrupts the cellular antioxidant defense system, thereby exacerbating lipid peroxide accumulation. Collectively, these data provide critical biochemical evidence that OMT-induced cell death in MKN28 cells is linked to the induction of lipid peroxidation, a core characteristic of ferroptosis.

The initiation and execution of ferroptosis are strictly dependent on the accumulation of labile iron ions within cells, which catalyze the Fenton reaction to generate reactive oxygen species (ROS) that propagate lipid peroxidation ^[6,7]^. Our iron ion detection assay revealed that OMT treatment significantly increased intracellular iron ion concentrations in MKN28 cells,. This iron overload likely creates a pro-oxidative intracellular microenvironment that amplifies lipid peroxidation cascades, thereby driving ferroptosis. The simultaneous induction of lipid peroxidation and iron accumulation by OMT—two non-redundant hallmarks of ferroptosis—strongly validates that OMT triggers this specific cell death pathway in MKN28 cells, distinguishing our findings from prior reports on OMT-induced apoptosis in other tumor types.

Ferroptosis is orchestrated by a complex network of signaling pathways and regulatory molecules, with the balance between pro-ferroptotic and anti-ferroptotic factors dictating cell fate^[8]^. ACSL4 (acyl-CoA synthetase long-chain family member 4) is a critical pro-ferroptotic mediator that selectively activates PUFAs, such as arachidonic acid and adrenic acid, for incorporation into cellular membranes, thereby increasing membrane susceptibility to peroxidative damage ^[9,10]^. In the current study, Western blot analysis demonstrated that OMT treatment significantly upregulated ACSL4 protein expression in MKN28 cells (Figure 4), suggesting that OMT may enhance lipid peroxidation by activating ACSL4-dependent PUFA metabolism. This finding aligns with recent studies showing that ACSL4 upregulation sensitizes gastric cancer cells to ferroptosis inducers, highlighting ACSL4 as a conserved mediator of ferroptosis in gastric cancer.

In contrast to ACSL4, GPX4 (glutathione peroxidase 4) and SLC7A11 (solute carrier family 7 member 11) function as core anti-ferroptotic regulators^[11,12]^. GPX4 is uniquely capable of reducing lipid hydroperoxides to non-toxic alcohols using GSH as a co-substrate, thereby preventing the propagation of lipid peroxidation. SLC7A11, a key component of the cystine-glutamate antiporter system Xc−, mediates the uptake of extracellular cystine, which is essential for intracellular GSH synthesis—an indispensable cofactor for GPX4 activity^[13-15]^. Our results showed that OMT treatment significantly downregulated the protein expressions of both GPX4 and SLC7A11 in MKN28 cells (Figure 4), leading to GSH depletion and impaired lipid peroxide clearance. This dual inhibition of the GPX4-GSH and SLC7A11-GSH antioxidant axes creates a synergistic pro-ferroptotic effect: SLC7A11 downregulation reduces GSH biosynthesis, while GPX4 suppression abolishes the cell’s capacity to neutralize existing lipid peroxides. This coordinated disruption of antioxidant defense mechanisms likely amplifies OMT-induced lipid peroxidation and iron accumulation, ultimately driving MKN28 cells toward ferroptosis. Our results showed that OMT significantly downregulated the protein expressions of GPX4 and SLC7A11, suggesting that OMT may promote lipid peroxidation and induce ferroptosis in gastric cancer MKN28 cells by inhibiting the GPX4-GSH and SLC7A11-GSH antioxidant systems.

This study makes a significant contribution to the field by identifying ferroptosis as a novel antitumor mechanism of OMT in gastric cancer, thereby providing a new theoretical basis and potential therapeutic target for gastric cancer treatment. OMT, as a natural product with low toxicity and favorable pharmacokinetic properties, holds promise for development as a ferroptosis-inducing agent for gastric cancer, particularly in cases where conventional therapies fail due to chemoresistance. However, several limitations of this work should be acknowledged to guide future research. Firstly, the current study was restricted to in vitro experiments using the MKN28 cell line, and the antitumor efficacy and ferroptosis-inducing capacity of OMT need to be validated in in vivo models to assess its translational potential. Secondly, gastric cancer is a highly heterogeneous disease, and the response to OMT may vary across different molecular subtypes. Future studies should evaluate OMT’s effect on a panel of gastric cancer cell lines with distinct genetic backgrounds to determine the generalizability of our findings. Thirdly, the upstream signaling pathways mediating OMT-induced regulation of ACSL4, GPX4, and SLC7A11 remain elusive. For example, the Nrf2 (nuclear factor erythroid 2-related factor 2) pathway, a master regulator of antioxidant responses, has been shown to transcriptionally upregulate SLC7A11 and GPX4; whether OMT modulates Nrf2 activity to downregulate these genes warrants further investigation. Additionally, the role of iron metabolism-related proteins (e.g., transferrin receptor 1, ferritin heavy chain 1) in OMT-induced iron accumulation requires detailed exploration to fully elucidate the molecular cascade underlying OMT-induced ferroptosis.

## 5. Conclusion

Oxymatrine can inhibit the viability of MKN28 cells in a dose- and time-dependent manner. It can induce an increase in intracellular lipid peroxidation level and iron ion concentration, while regulating the expressions of ferroptosis-related genes and proteins, indicating that oxymatrine can induce ferroptosis in gastric cancer MKN28 cells. This study provides a new theoretical foundation and potential therapeutic direction for the application of oxymatrine in gastric cancer treatment.

## Declarations

### Ethics approval and consent to participate

All animal experiments in this study were reviewed and approved by the Animal Ethics Committee of Ningxia Medical University

